# Glycolytic reprogramming shapes the epigenetic landscape of activated CD4^+^ T Cells in Juvenile Idiopathic Arthritis

**DOI:** 10.1101/2024.02.14.580242

**Authors:** Enric Mocholi, Theo Chalkiadakis, Can Gulersonmez, Edwin Stigter, Bas Vastert, Jorg van Loosdregt, Stefan Prekovic, Paul J Coffer

**Affiliations:** Center for Molecular Medicine, University Medical Center Utrecht, The Netherlands; Regenerative Medicine Center, University Medical Center Utrecht, The Netherlands; Center for Translational Immunology, University Medical Center Utrecht, The Netherlands; Division of Pediatrics, University Medical Center Utrecht, The Netherlands

## Abstract

Juvenile Idiopathic Arthritis (JIA) describes a heterogeneous group of autoimmune conditions with an unknown cause and childhood onset. It is characterized by the accumulation of mononuclear cells, notably activated CD4^+^ memory/effector T (Tmem/Teff) cells, within the synovial fluid of affected joints. JIA CD4^+^ T cells exhibit a unique epigenomic signature linked to inflammation, however, the molecular mechanisms driving this remain unclear. Here we show that CD4^+^ T cells isolated from JIA synovial fluid (SF) exhibit abnormal intracellular metabolism marked by heightened glycolysis after activation driving transcriptional reprogramming. Epigenetic profiling between activated healthy controls and JIA patients allowed the definition of specific disease-related enhancers upregulated in SF-derived JIA CD4^+^ T cells. Pharmacological inhibition of glycolytic flux affected the expression of genes associated with these enhancers. When activated in the presence of JIA SF, CD4^+^ T cells obtained from healthy control (HC) subjects, displayed heightened glycolytic activity compared to paired plasma. Moreover, this also led to increased H3K27ac at JIA-specific genes. Increased H3K27ac was dependent on glycolytic flux, but not oxidative phosphorylation. Inhibition of glycolysis also specifically affected the transcription of genes upregulated during T cell activation in the presence of SF. Inhibiting the glycolytic enzyme pyruvate dehydrogenase (PDH) reduced JIA-associated gene expression. Taken together, these findings demonstrate that for JIA, the inflammatory microenvironment can modulate T cell activation-driven transcriptional programs through a glycolysis-mediated pathway. Specific targeting of this T cell metabolism-epigenetic axis may provide avenues for intervention during the development of autoinflammatory disease.

## Introduction

Juvenile idiopathic arthritis (JIA) is a heterogeneous group of autoimmune conditions, characterized by chronic arthritis with an unknown cause and onset before the age of 16 years. It is marked by the accumulation of mononuclear cell populations, notably including activated CD4^+^ memory/effector T (Tmem/Teff) cells, within the synovial fluid of afflicted joints (Prakken et al., 2011). The pivotal role of Tmem/Teff cells in the pathophysiology of JIA has been well-established^1^. JIA is a relevant model for the exploration of autoimmune pathogenesis, primarily due to its early onset and the potential for in-depth analysis of immune cells derived from inflamed anatomical sites. Previously, we identified a unique disease-specific epigenomic signature linked to inflammation in CD4^+^ T cells isolated from the synovial fluid of JIA patients^2^. However, the molecular mechanisms underlying epigenome dysregulation in JIA T cells, or T cells associated with other autoinflammatory conditions, remain unclear.

Intracellular metabolism is critical in modulating immune activation influencing fundamental processes such as proliferation, differentiation, cell fate, and functions^3^. Upon activation of the T-cell receptor (TCR), a metabolic program is initiated that not only enhances mitochondrial function but also engages aerobic glycolysis^3^. Recent studies have identified an essential role for glycolysis in orchestrating epigenetic and transcriptional changes during T-cell activation and differentiation^4–7^. However, it remains unclear how this is impacted when considering inflammatory environments. Initial investigations in individuals with rheumatoid arthritis (RA) have delineated a disease-specific metabolic signature^8^. This underlies the transformation of naive CD4^+^ T cells into pro-inflammatory helper T cells, which subsequently infiltrate joints and provoke inflammation through immunogenic cell death^9^. These pro-inflammatory T cells infiltrate the RA synovium, contributing to tissue damage^10^. Activated RA T cells are unable to upregulate expression of the key glycolytic enzyme 6-phosphofructo-2-kinase/fructose-2,6-bisphosphatase 3 (PFKFB3), an indication that patient-derived cells utilize glucose differently (Yang et al., 2013). In a human tissue-mouse chimera model, selective knockdown of PFKFB3 in adoptively transferred healthy T cells was sufficient to induce robust synovial inflammation, generating a transcriptome pattern reminiscent of rheumatoid synovitis^11^. Conversely, rescue of PFKFB3 expression in adoptively transferred RA T cells was effective in dampening synovial tissue inflammation. In essence, by shifting the ratio of PFKFB3/G6PD (glucose-6-phosphate-deshydrogenase), RA T cells divert glucose from ATP production toward biosynthesis, generating inflammation-inducing effector cells. Furthermore, mitochondrial oxygen consumption is significantly suppressed in RA T cells despite mitochondrial mass being comparable to that in healthy T cells^12^. A better understanding of the role of metabolic signals in T cell specification opens the possibility for immunomodulation before the end-stage of synovial inflammation encountered in clinical practice^8^.

Here, we have examined the interplay between metabolic reprogramming and epigenetic regulation in controlling CD4^+^ T cell activation in JIA. T cells from the synovial fluid of individuals with JIA display anomalous intracellular metabolism and epigenetic profiles. Inhibiting glycolytic flux affected the expression of genes associated with this aberrant epigenetic profile. Furthermore, synovial fluid has the capacity to activate acetylation of enhancers normally associated with pathogenic JIA T cells. These changes in the epigenetic profile are contingent upon glycolysis-derived acetyl-CoA production facilitated by pyruvate dehydrogenase. This underscores the close connection between glycolysis, transcriptional responses, and T cell functionality. Understanding the impact of metabolic changes in pathological settings provides a foundation for developing therapeutic interventions aimed at selectively targeting these pathways.

## Materials and Methods

### Collection of SF and PB Patient Samples

Peripheral blood (PB) was obtained from healthy donors (HC) under the Minidonor Dienst Program (UMC Hospital). Twenty-four oligoarticular Juvenile Idiopathic Arthritis (JIA) patients were included in this study who at the time of sampling all had active disease and underwent therapeutic joint aspiration allowing SF collection. PB was drawn at the same moment via vein puncture or intravenous drip. Informed consent was obtained from all patients either directly or from parents/guardians when the patients were younger than age 12 years. The study procedures were approved by the Institutional Review Board of the University Medical Center Utrecht (UMCU; METC nr: 11-499c) and performed according to the principles expressed in the Helsinki Declaration. Hyaluronic acid was broken down in SF samples by 30 min incubation at 37°C with Hyaluronidase (Sigma). Synovial fluid mononuclear cells (SFMCs) and peripheral blood mononuclear cells (PBMCs) were isolated using Ficoll Isopaque density gradient centrifugation (GE Healthcare Bio-Sciences AB) and were used fresh or after freezing in FCS (Invitrogen) containing 10% DMSO (Sigma-Aldrich).

### Cell isolation and culture

CD4+ T cells were isolated from SFMCs and PBMCs using MagniSortTM Human CD4+ T cell Enrichment Kit (eBioscience 8804-6811-74). The Cd4 T cells were cultured always in RPMI Medium 1640 + GlutaMAX supplemented with 100 U/ml penicillin, 100 mg/ml streptomycin (all obtained from Life Technologies), and 10% heat-inactivated human AB-positive serum (Invitrogen) at 37°C in 5% CO2. Where indicated, CD4 T cells were activated with 0.5 µg/ml plate-bound anti-CD3 (eBioscience; 16-0037) and 0.5 µg/ml anti-CD28 (eBioscience; 16-0289-85) during 12, 24 or 48 hours. Where indicated cells were treated with Oligomycin (1µM Sigma Aldrich), 2DG (50mM Sigma Aldrich), 6,8-Bis(benzylthio) octanoic acid (Sigma Aldrich), or with 30% Synovial Fluid (SF) from JIA Patients.

### Chromatin-immunoprecipitation (ChIP)

For each sample, cells were crosslinked with 2% formaldehyde and crosslinking was stopped by adding 0.2 M glycine. Nuclei were isolated in 50 mM Tris (pH 7.5), 150 mM NaCl, 5 mM EDTA, 0.5% NP-40, and 1% Triton X-100 and lysed in 20 mM Tris (pH 7.5), 150 mM NaCl, 2 mM EDTA, 1% NP-40, 0.3% SDS. Lysates were sheared using Covaris microTUBE (duty cycle 20%, intensity 3, 200 cycles per burst, 60-s cycle time, eight cycles) and diluted in 20 mM Tris (pH 8.0), 150 mM NaCl, 2 mM EDTA, 1% X-100. Sheared DNA was incubated overnight with anti-histone H3 acetyl K27 antibody (ab4729; Abcam) pre-coupled to protein A/G magnetic beads. Beads were washed and crosslinking was reversed by adding 1% SDS, 100 mM NaHCO3, 200 mM NaCl, and 300 mg/ml proteinase K. DNA was purified using ChIP DNA Clean & Concentrator kit (Zymo Research). The Chip DNA was used for sequencing or qPCR.

### DNA-Sequencing

End repair, a-tailing, and ligation of sequence adaptors were done using Truseq nano DNA sample preparation kit (Illumina). Samples were PCR amplified, checked for the proper size range, and for the absence of adaptor dimers on a 2% agarose gel, and barcoded libraries were sequenced 75 bp single-end on Illumina NextSeq500 sequencer (Utrecht DNA sequencing facility).

### ChIP-qPCR

Real-time PCR was performed with PowerSYBR (Applied Biosystems) using a StepOnePlus Real-Time-PCR system (Applied Biosystems). The expression of each gene was normalized to a negative region. All the primers for the Chip-qPCR they were designed based on or Chip-seq: qPCR CHIP negative region: F-GAGCCAGGGTTTCTCTGATTC, R-CCTCAGTGATCAGCCCTAAATG; qPCR CHIP IL10RA: F-CCCGCTCCATTAAAGTTCTCC, R-TTTCAGCCTCTTCCACTTCC; qPCR CHIP TRPS1: F-AGTTATTTGGAGGGACAGCG, R-GTGATTAAATGCCTGACAGCG; qPCR CHIP PNRC1: F-GCTGTTTCACTTTCTCCCTTTG, R-AGGATTGTGAGACTTTGGGATC; qPCR CHIP CD74: F-CACTTACCAAGCTCTCCTTCG, R-CTACTTTCTGGAGGTGTGATCC and qPCR CHIP CD80: F-TCTACTCCCACCTCTGAATCC, R-CTAAAGTCTTCCTCATCCCACC.

### Real-Time PCR

RNA was isolated from cells using RNasy Mini Kit (Qiagen) and cDNA was synthesized using Superscript-III First-Strand Synthesis System (Life Technologies). Real-time PCR was performed with PowerSYBR (Applied Biosystems) using a StepOnePlus Real-Time-PCR system (Applied Biosystems). Expression of each gene was normalized to β2M. The following primer sets were used: RNA q-PCR β2M: F-ATGAGTATGCCTGGCCGTGTGA, R-GGCATCTTCAAACCTCCATG; RNA qPCR IL10RA: F-AAGTGGCGCTCCTGAGGTAT, R-GCTGTCTGTGCTATTGCTGC; RNA qPCR TRPS1: F-ACCAGCATGCAGAGTAATATGGT, R-GTTTCCTCCCTTACTGGGGC; RNA qPCR PNRC1: F-CCCCCTCAGGAAAGAGGTTTT, R-TGCCATCAGCTCCCTGTTTT; RNA qPCR CD74: F-TGGCCTTCTGTGGACGAATC, R-CAGTGACTCTGGAGCAGGTG; RNA qPCR CD80: F-CCGAGTACAAGAACCGGACC, R-GGTGTAGGGAAGTCAGCTTTGA.

### Seahorse assays

T cells were stimulated with anti-CD3 and anti-CD28 for 12, 24, and 48 hours. Oxygen consumption rates (OCR) and extracellular acidification rates (ECAR) were measured in XF media (non-buffered RPMI 1640 containing 10 mM glucose, 2 mM L-glutamine, and 1 mM sodium pyruvate) under basal conditions and in response to glucose 30mM, 1uM oligomycin, and 50mM of 2DG, on an XF-24 Extracellular Flux Analyzers (Seahorse Bioscience).

### Measurements of Acetyl-CoA Levels

The intracellular levels of acetyl-CoA were detected by using an Acetyl-CoA Assay Kit (Biovision, Milpitas CA), following the manufacturer’s instructions.

### Sample Preparation and LC-MS Measurement of glycolytic/TCA cycle intermediates

Polar metabolites were extracted from 2×10e6 T-cells isolated from peripheral blood or synovial fluid by adding 500µL ice-cold methanol solution containing internal standards. The cell lysis was performed by using a bullet blender and 0.5 mm glass beads (NextAdvance, USA) in the sample tube. After cell lysis the extracts were centrifuged for 10min at 17000xg and 4°C. The complete supernatant was transferred into a fresh 1.5mL tube and evaporated to dryness in a vacuum concentrator (Labconco CentriVap, USA) for 1h. The dried metabolite pellet was dissolved in 120µL of water/acetonitrile solution (95%/5%, v/v) prior to transfer the material into LC glass vials.

The chromatographic separation was performed using a ThermoFischer Accela UHPLC System equipped with a Waters Acquity C8 column (2.1 x 150 mm, 1.8 µm) the outlet of which was connected to an LTQ-Orbitrap XL MS System (ThermoFisher Scientific, USA and Waters, USA). The LC flow rate was set to 150µL/min and the column oven temperature was adjusted to 35°C. Five microliters of the sample was injected on column. The weak and the strong eluents were 100% Milli-Q water and methanol/Milli-Q water (95%/5%, v/v) respectively and both solvents contain 6.5mM ammonium bicarbonate at pH 8.0. The strong eluent starts at 0% for 1min and increases over 1-5min from 0% to 70% and from 70% to 98% between 5-5.5min following an isocratic run over 5.5-15.5min and a drop back to the initial 0% composition between 15.5-22min. The MS data were acquired in full scan mode between 55-900 amu. The peak picking and integration was performed with Thermo Xcalibur Software (v2.4). The target metabolites were identified by running authenticated standards in parallel and comparing their retention time and m/z values. The obtained peak areas were corrected by the peak area of the corresponding internal standard.

### Statistical Analysis

For ChIP-seq and RNA-seq analysis, p values were adjusted with the Benjamini-Hochberg procedure. For ChIP-seq regions with a significantly different H3K27ac signal were defined using a false discovery rate (FDR) <0.05.

## Results

### SF JIA T cells are metabolically reprogrammed with increased activation-induced glycolysis

To evaluate the effect of local inflammation on CD4^+^ T cell metabolic responses, we conducted an analysis of intermediate metabolites associated with glycolysis and the Tricarboxylic Acid (TCA) cycle. Specifically, we compared the levels of these metabolites in CD4^+^ T cells derived from the Synovial Fluid (SF) of patients with active Juvenile Idiopathic Arthritis (JIA) to those from healthy controls obtained from peripheral blood (PB HC CD4^+^). CD4^+^ T cells were polyclonally activated with anti-CD3/CD28 in the presence of D-Glucose-^13^C_6_. We subsequently measured the levels of intracellular ^13^C-labeled intermediate metabolites associated with glycolysis and TCA by LC-MS mass spectrometry (**Figure 1a**). Increased concentrations of intermediary metabolites linked to glycolysis and the TCA cycle in SF JIA T cells and PB HC T cells were observed.

**Figure 1.**
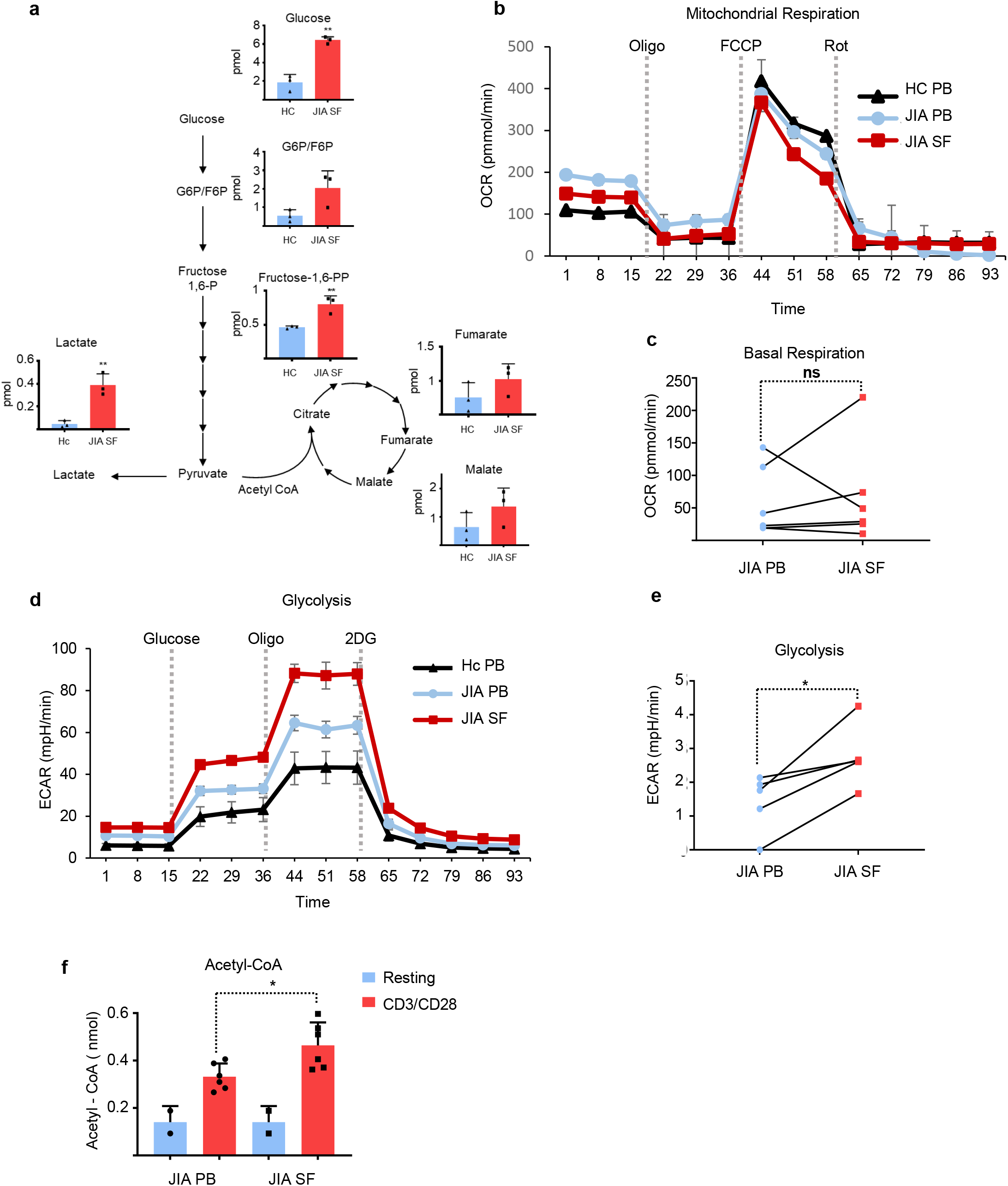
Enhanced activation-induced glycolysis in SF JIA CD4^+^ T Cells. **(a)** HC PB or JIA SF CD4^+^ T cells were activated for 24h with anti-CD3/CD28. Cell lysates from 3 biological replicates were extracted and analyzed using high-resolution LC-QE-MS to determine cellular metabolites. Representation of relative levels of metabolites in the glycolysis and TCA-cycle pathways are shown. Data are shown as mean ± SD of triplicate samples. **(b-c)** JIA from PB or SF CD4^+^ T cells were activated for 24h with CD3/CD28 and extracellular acidification rates (OCR) were measured by Seahorse technology. **(d-e)** JIA from PB or SF CD4^+^ T cells were activated for 24h with CD3/CD28 and extracellular acidification rates (ECAR) were measured by Seahorse technology. **(f)** HC PB or JIA SF CD4^+^ T cells isolated were activated for 24h with anti-CD3/CD28 and acetyl-CoA levels were measured. All graphs represent mean +/-SD. One-way ANOVA or Paired Student’s T-test measured statistical significance. * P<0.05; **P<0.01.

To further investigate the glycolytic responses of PB HC CD4^+^, PB JIA CD4^+^, and SF JIA CD4^+^ T cells, we performed *in vitro* activation using anti-CD3/CD28 for 24 hours and assessed mitochondrial respiration and anaerobic glycolysis through measurement of Oxygen Consumption Rate (OCR) and Extracellular Acidification Rate (ECAR), respectively. We did not observe significant differences in mitochondrial respiration (**Figure 1b**), ATP production by mitochondria (**Figure S1a-b**, or maximal respiration levels (**Figure 1c**). However, basal glycolysis was markedly increased in JIA SF CD4^+^ T cells following T cell activation, compared to both healthy controls and CD4^+^ T cells isolated from the peripheral blood of the same JIA patients (**Figure 1d-e**). These cells also exhibited changes in glycolytic capacity and glycolytic reserve (**Figure S1c-d**), but these differences were not statistically significant.

Glycolytic flux can directly reprogram the epigenetic landscape as exemplified, for example, by increased histone acetylation^13^. We measured the intracellular levels of acetyl-CoA in JIA CD4^+^ T cells isolated from both the peripheral blood and synovial fluid either in a resting state or stimulated with anti-CD3/CD28 for 24 hours. CD3/CD28-mediated activation induced an increase in intracellular acetyl-CoA levels, but the increase was significantly higher in JIA SF CD4^+^ T cells (**Figure 1f**).

Taken together, SF JIA T cells exhibit changes in intracellular metabolism marked by elevated activation-induced levels of TCA and glycolytic intermediate metabolites. This metabolic dysregulation is concomitant with increased glycolytic flux and acetyl-CoA production upon activation.

### JIA CD4^+^ T cells exhibit an abnormal activation-induced H3K27ac landscape

To gain further insights into epigenetic and transcriptomic reprogramming occurring during the activation of SF JIA CD4^+^ T cells, we conducted chromatin immunoprecipitation followed by DNA sequencing (ChIP-seq) experiments targeting histone 3 Lysine 27 acetylation (H3K27ac) a histone modification associated with activated promoters and enhancers^14^. To this end, we isolated CD4^+^ T cells from both PB HC and SF JIA patients. These cells were cultured *in vitro* under resting conditions for 24 hours to minimize the direct impact of the synovial inflammatory milieu. Subsequently, cells were stimulated with anti-CD3/CD28 for 24 hours (**Figure 2a**). Significant differences were observed in the H3K27ac profiles of CD4^+^ T cells isolated from SF JIA compared to those from healthy controls. JIA T cells exhibited over 6000 regions with increased acetylation, while more than 5000 regions lost acetylation compared to the healthy control group. Further evaluation using Gene Ontology (GO)-term analysis highlighted that regions that were more acetylated on the JIA CD4^+^ T cells were associated with genes involved in T cell activation, differentiation, proliferation, and cytokine production (**Figure 2b, S2a**). These findings demonstrate distinct activation-induced H3K27ac profiles in SF JIA T cells compared to PB HC. Additionally, using gene set enrichment analysis (GSEA) we observed a robust correlation between genes exhibiting increased H3K27 acetylation in JIA T cells and those undergoing upregulation upon T cell activation (**Figure S2b**). Furthermore, there was a significant association between increased H3K27 acetylation (within 5 kb of the transcription start site (TSS)) in SF JIA T cells and genes linked to crucial immune effector pathways such as interferon gamma response, TNF-α signaling via NF-KB, and IL-2-STAT5 signaling (**Figure 2c**). Approximately one-third of the annotated peaks, upregulated in SF JIA T cells compared to HC T cells, were found to localize to promoter regions (**Figure 2d**). This observation was further supported by analysis of transcription factor binding sites, where >25% are in proximity (<5kb) to H3K27ac peaks (**Figure 2e**). The most statistically significant predicted transcription factor binding sites were for ETS1, KLF6, RUNX1, AP1, and PLAGL1 (**Figure 2f**).

**Figure 2.**
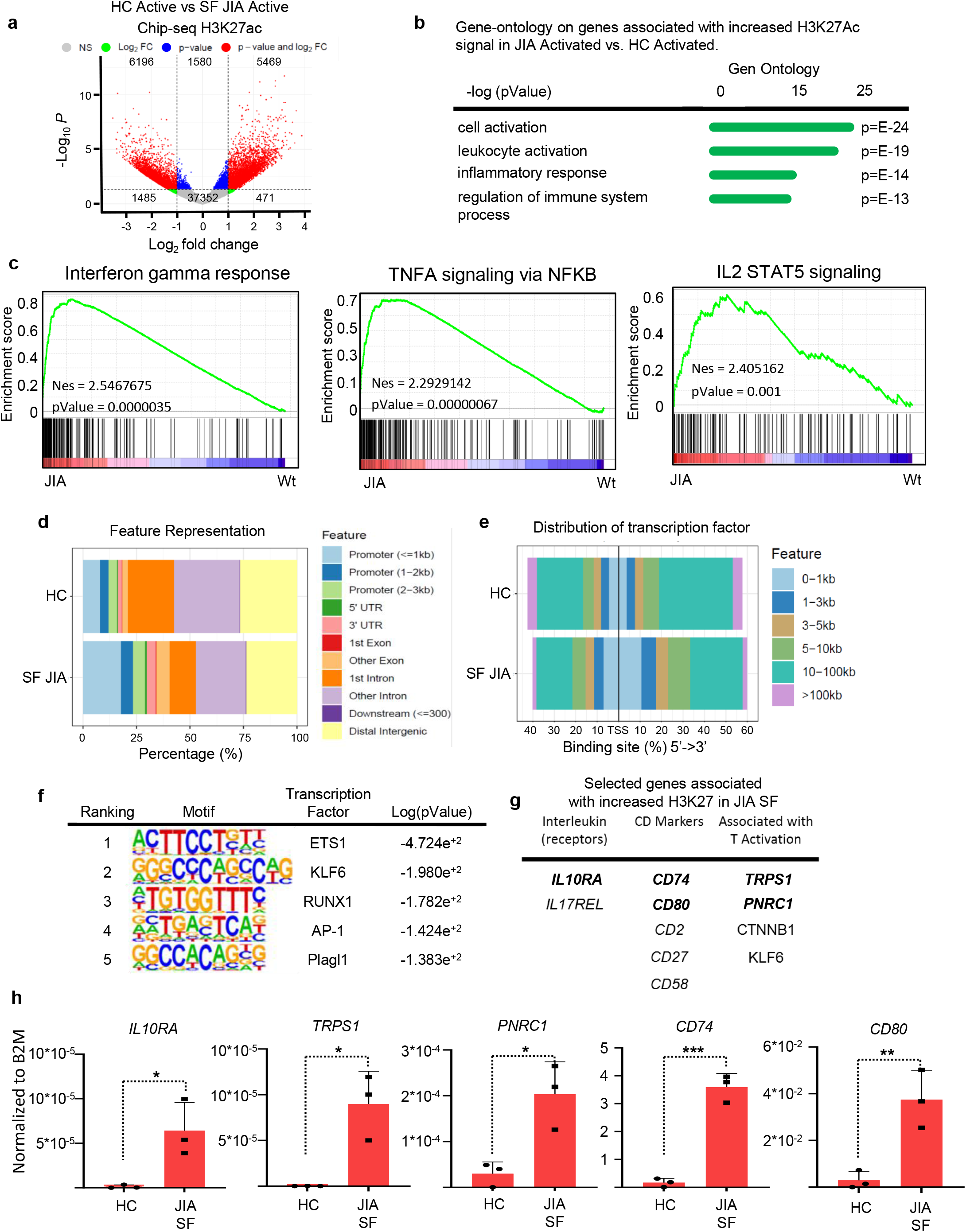
JIA CD4^+^ T cells exhibit an altered H3K27ac landscape. **(a)** Volcano plot of H3K27ac signal, based on comparisons of all replicates within the depicted groups. Red dots indicate enhancers with an FDR <0.05. **(b)** Gene ontology terms, ranked by enrichment scores genes that are up regulated on JIA activated versus HC activated CD4^+^ T cells. (**c)** Gene set enrichment analysis between differentially expressed genes from JIA activated CD4^+^ cells and hallmarks of Interferon-gamma response, TNFA signaling via NFKB and IL2 STAT5 signaling. **(d)** Genomic distribution of H3K27ac peaks identified in ChIP-Seq data. **(e)** Distribution of transcription factor-binding loci relative to 5’ ends of genes. **(f)** Transcription factor binding motifs enriched in SF JIA T cells. **(g)** Selected genes upregulated in activated JIA SF-derived CD4+ T cells. **(h)** qRT-PCR measured mRNA expression of *IL10RA*, *TRPS1*, *PNRC1*, *CD74*, *CD80* in human CD4+ T from HC PB or JIA SF cells activated 24h with anti-CD3/CD28. All graphs represent mean +/-SD. One-way ANOVA or Student’s T-test measured statistical significance. * P<0.05; **P<0.01; ***P<0.001; ****P<0.0001.

Through a comprehensive comparison of H3K27ac datasets between healthy controls and SF JIA, we successfully identified specific disease-related promotors that exhibited upregulation in SF JIA CD4^+^ T cells in contrast to PB HC T cells (**Figure 2g, S2c-d**). To validate the relevance of these H3K27ac regions, we measured the expression levels of *IL10RA*, *TRPS1*, *PNRC1*, *CD74*, and *CD80* upon *in vitro* activation of SF JIA T cells. We indeed observed significant upregulation of these genes in SF JIA T cells compared to PB HC T cells (**Figure 2h)**. These data substantiate the correlation between the aberrant epigenome of these cells and their transcriptome.

Collectively, these results indicate that T cells isolated from the SF of JIA patients maintain an aberrant epigenome even upon ex-*vivo* activation. Notably, there is an increased accessibility in the promoter regions of multiple genes involved in immune effector pathways.

### JIA synovial fluid drives metabolic reprogramming and a dysregulated H3K27ac landscape

To further investigate the potential link between the metabolic dysregulation observed in SF JIA CD4^+^ T cells and their aberrant H3K27ac profile, HC CD4^+^ T cells were activated *in vitro* with anti-CD3/CD28 for 24 hours in the presence of either 30% SF or 30% plasma from JIA patients. Using Seahorse metabolic profiling, we evaluated changes in OCR and ECAR as indicators of mitochondrial respiration and anaerobic glycolysis, respectively. Synovial fluid did not significantly impact the oxygen consumption rate of T cells compared to plasma from the same patients (**Figure 3a-b, S3 a-b**). However, CD4^+^ T cells stimulated in the presence of SF displayed a significant elevation in glycolysis, glycolytic reserve, and glycolytic capacity, compared to those activated in the presence of plasma derived from the same individuals (**Figure 3c-d, S3c-d**). These findings resemble the data of SF-activated CD4^+^ T cells (**Figure 1d-e**), underscoring the distinctive metabolic profile of JIA SF, and suggesting a potential role for SF in enhancing glycolytic pathways and fueling their effector functions.

**Figure 3.**
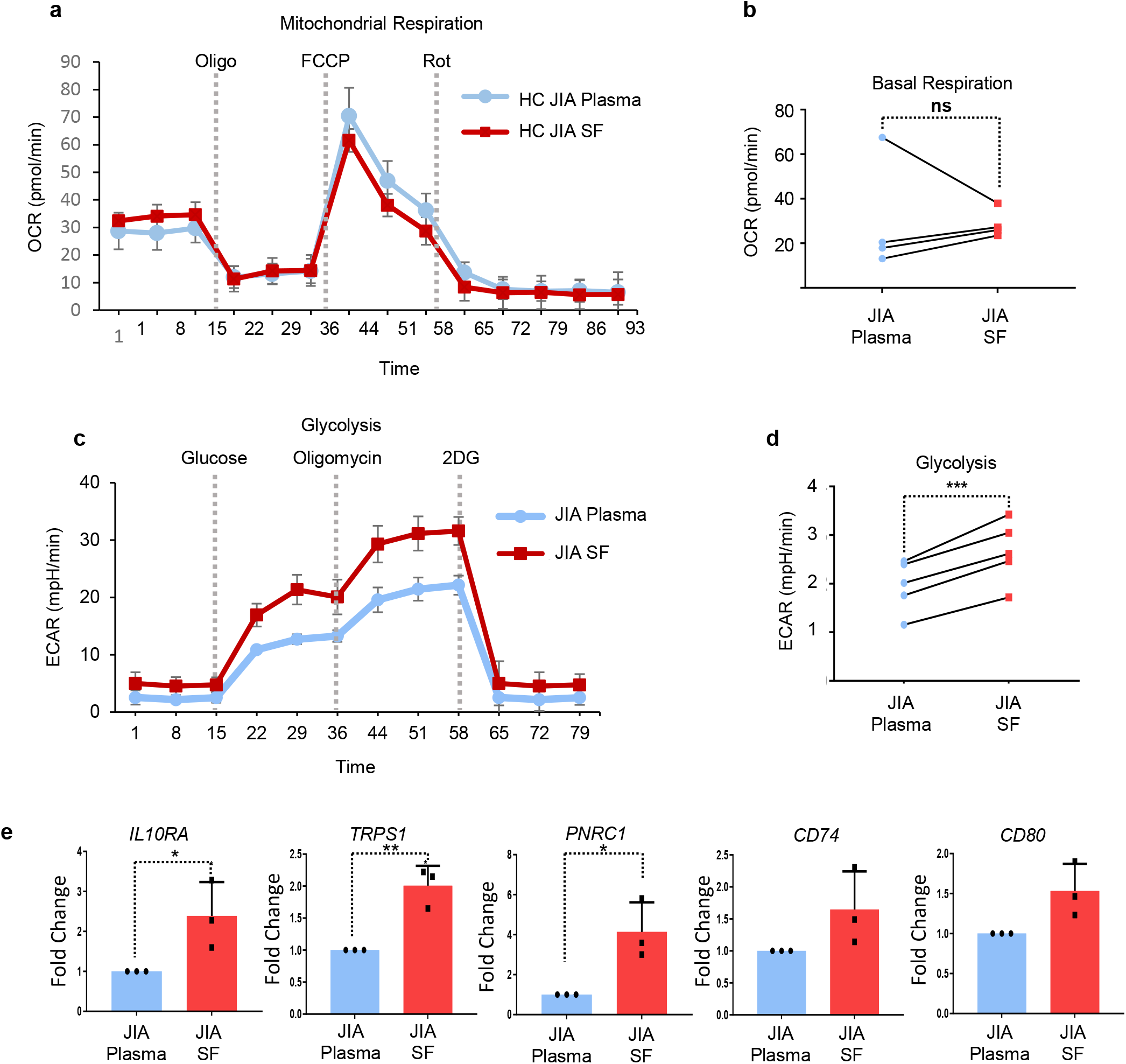
JIA synovial fluid promotes metabolic reprogramming and H3K27ac landscape remodeling. (**a-b)** HC PB CD4^+^ T cells were activated for 20h with anti-CD3/CD28 in presence of JIA SF or plasma (30%) and extracellular acidification rates (OCR) measured by Seahorse technology. **(c-d)** HC PB CD4^+^ T cells were activated for 20h with anti-CD3/CD28 in the presence of JIA SF or plasma (30%) and extracellular acidification rates (ECAR) measured by Seahorse technology. **(e)** ChIP-qPCR of *IL10R*, *TRPS1*, *PNRC1*, *CD74* and *CD80* promotor regions from human healthy CD4^+^ T cells activated with anti-CD3/CD28 in the presence of JIA SF or plasma (30%). All graphs represent mean +/-SD. Student’s T-test measured statistical significance. * P<0.05; **P<0.01.

Subsequently, we investigated the effects of SF on HC CD4^+^ T cells, focusing on H3K27ac modification at enhancers specifically upregulated in JIA SF CD4^+^ T cells. PB HC CD4+ T cells were activated with anti-CD3/CD28 for 24 hours, in the presence of either 30% SF or 30% plasma obtained from JIA patients. Employing H3K27ac ChIP-qPCR analysis, we quantified the acetylation levels of enhancers associated with *IL10RA*, *TRPS1*, *PNRC1*, *CD74*, and *CD80* genes (**Figure 3e**). Notably, the activation of CD4^+^ T cells in the presence of SF significantly increased H3K27ac at these JIA-associated promotors compared to matched plasma. Taken together, these data suggest that SF can impact both the rate of glycolysis and the epigenomic landscape of CD4^+^ T cells, thereby contributing to the upregulation of disease-related H3K27ac-mediated chromatin remodeling.

### Glycolytic control of H3K27ac-mediated chromatin remodeling in JIA CD4^+^ T cells

To further evaluate the impact of glycolysis on H3K27ac changes during CD4^+^ T cell activation, we activated JIA SF CD4^+^ T cells for 24-hours with or without 2-deoxy-D-glucose (2DG), an inhibitor of glucose-6-phosphate production which thereby inhibits glycolysis. Employing ChIP-seq analysis targeting H3K27ac, we observed that inhibiting glycolysis prevented epigenome remodeling after CD4^+^ T cell activation (**Figure 4a**).

**Figure 4.**
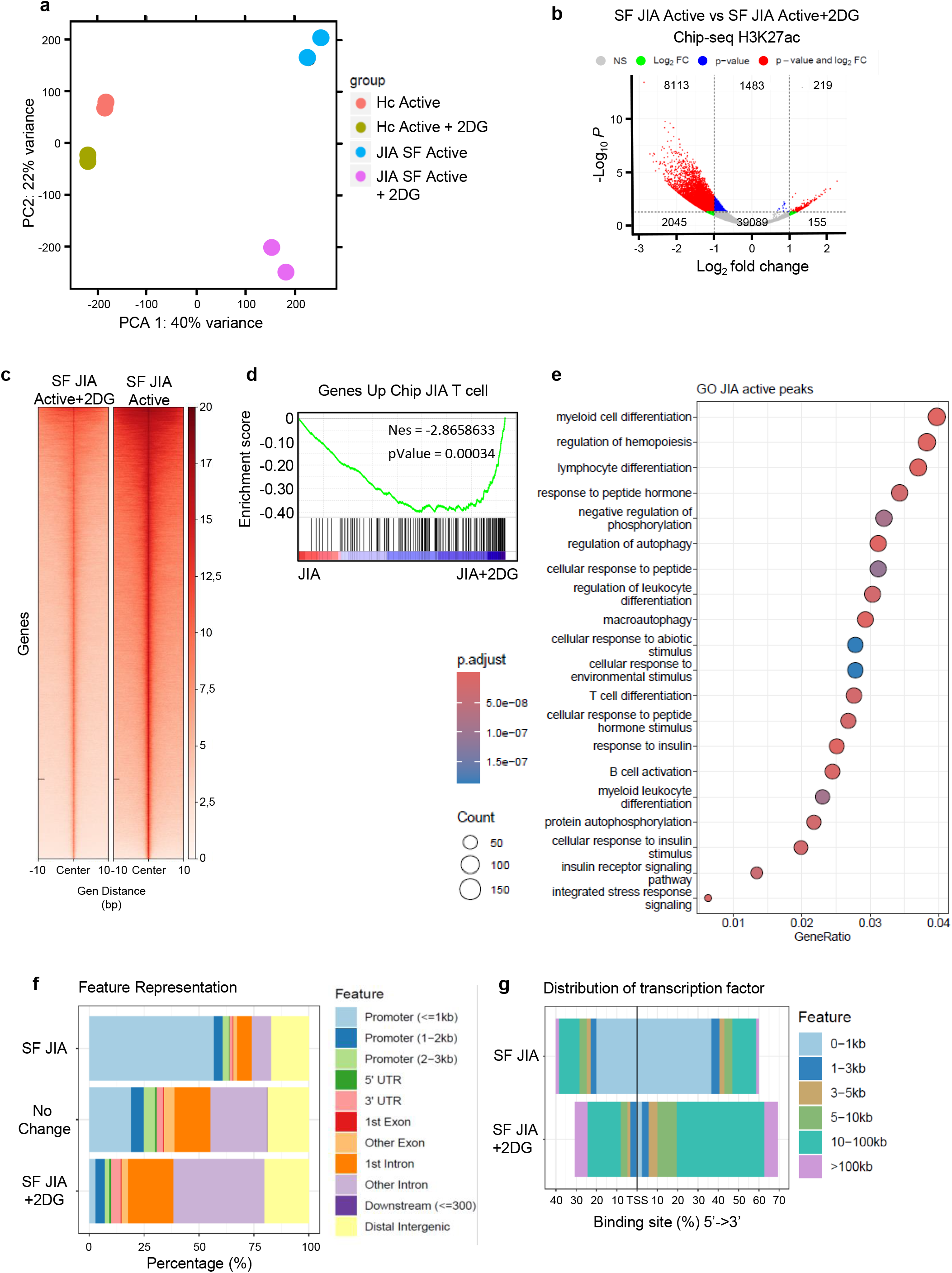
Glycolytic modulation of H3K27ac-driven chromatin remodeling in JIA CD4^+^ T Cells. **(a)** Principal component analysis based on H3K27ac signal. **(b)** Volcano plots of H3K27ac signal, based on comparisons of all replicates within the depicted groups. Red dots indicate enhancers with an FDR <0.05. **(c)** Plot of the percentages of significantly decreased H3K27ac peaks at enhancers and promoters on the JIA T cells activated in the presence of 2DG. **(d)** Gene set enrichment analysis comparing the H3K27ac signal in the T cell gene set of JIA CD4^+^ T cells to the Chip-seq data of JIA T CD4^+^ cells treated with 2DG. **(e)** Gene ontology terms are ranked by enrichment scores of genes (Es) that are up-regulated on JIA-activated CD4^+^ T cells versus activated in the presence of 2DG. **(f)** Genomic distribution of peaks identified in ChIP-Seq data. **(g)** Distribution of transcription factor-binding loci relative to 5’ ends of genes.

The inhibition of glycolysis markedly decreased histone acetylation after T cell activation in JIA SF T cells (**Figure 4b-c**). Genes affected by inhibition of glycolysis in SF-derived JIA T cells were found to correlate with the genes upregulated during T cell activation in SF compared to PB healthy T cells (**Figure 4d**). Moreover, inhibition of glycolysis significantly affected genes associated with CD4^+^ T cell activation and function (**Figure 4e**). The majority of peaks affected by 2DG-treatment in SF JIA T cells are annotated as promoter regions (**Figure 4f**), In contrast, HC T cells exhibit a distinct pattern, with only approximately 25% of the regions affected by glycolysis inhibition during activation being identified as promoter regions (**Figure S4a**). This observation was supported by an examination of transcription factor binding sites associated with these peaks (**Figure 4g, S4b**). Again, these results show that the impact of inhibiting glycolysis on promoter regions was more pronounced in SF JIA T cells compared to PB T cells (**Figure S4a**). These findings support a role for glycolysis in driving dysregulation of H3K27ac-associated promoter regions during activation in SF from JIA patients.

### Essential role of pyruvate-derived acetyl-CoA in mediating SF-driven changes in H3K27ac

To gain further insight into the role of glycolysis in the activation of disease-related enhancers, HC CD4^+^ T cells were activated for 24 hours with anti-CD3/CD28 in the presence of either plasma or JIA SF, with the addition of 2DG or oligomycin. Activation of CD4^+^ T cells in the presence of JIA SF resulted in a significant increase in H3K27ac levels at JIA SF-specific enhancers, compared to cells activated with plasma alone (**Figure 5a**). Importantly, this increase in H3K27ac was found to be dependent on glycolytic flux, rather than OXPHOS, as shown by the inhibition observed with 2DG but not oligomycin. To further validate these findings, gene expression was evaluated by quantitative RT-PCR (qRT-PCR). Increased H3K27ac levels were indeed associated with elevated gene expression. Furthermore, the inhibition of glycolysis had a more pronounced impact on gene expression compared to the inhibition of OXPHOS (**Figure 5b**).

**Figure 5.**
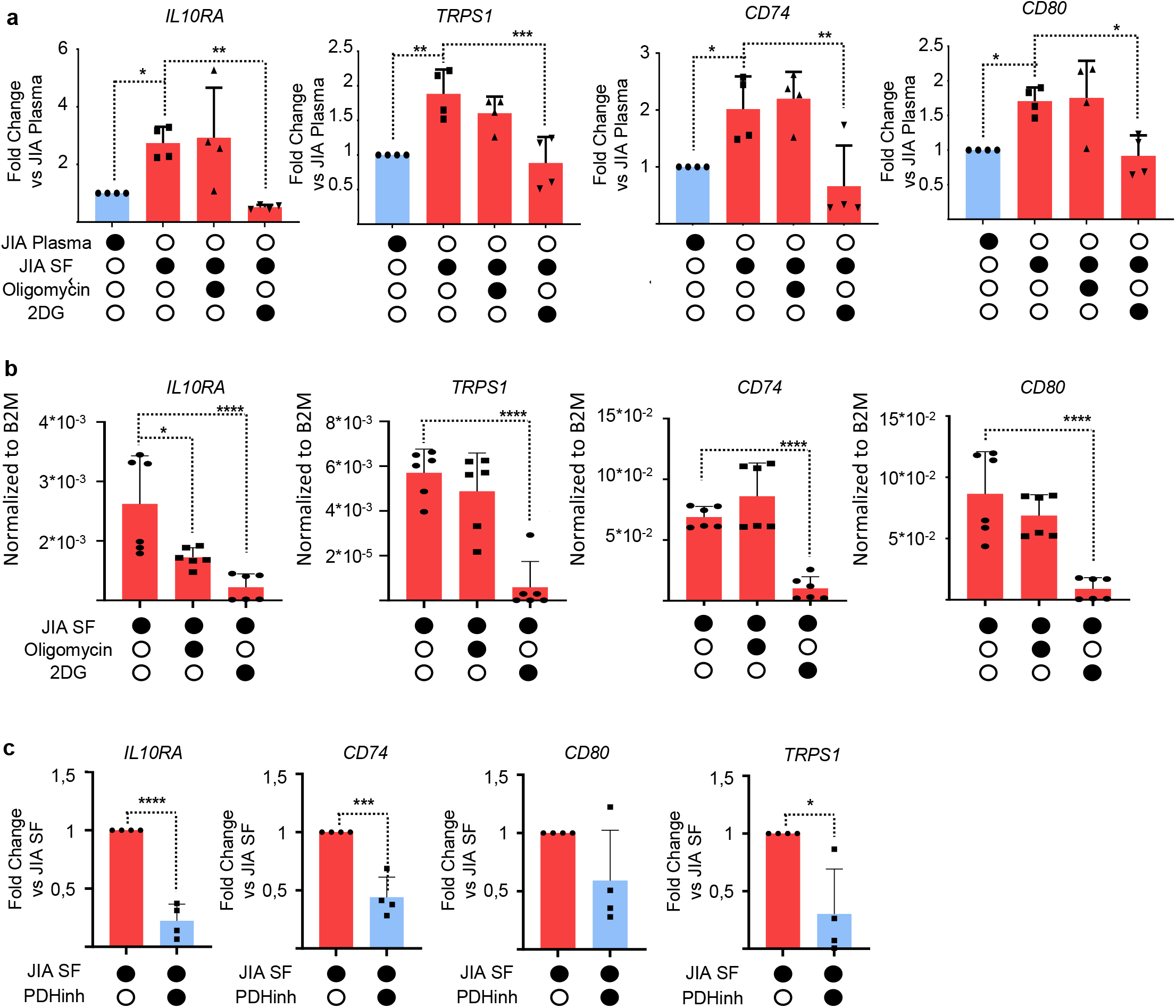
Glycolysis-driven pyruvate-metabolism is required for SF-mediated changes in H3K27ac. **(a)** ChIP-qPCR of *IL10R*, *TRPS1*, *PNRC1*, *CD74*, and *CD80* promotor regions from human CD4^+^ T cells activated with anti-CD3/CD28 in the presence of JIA SF or plasma (30%), and in the presence or absence of oligomycin or 2DG for 20h. **(b)** mRNA expression of *IL10RA*, *TRPS1*, *PNRC1*, *CD74*, and *CD80* was measured by qRT-PCR in human CD4^+^ T from HC PB in the presence of JIA SF, oligomycin or 2DG cells activated 24h with anti-CD3/CD28. **(c)** mRNA expression of *IL10RA*, *TRPS1*, *CD74*, and *CD80* was measured by qRT-PCR in human CD4^+^ T from HC PB in presence of JIA SF and 6,8-Bis(benzylthio) octanoic acid (PDH-inhibitor) cells activated 24h with anti-CD3/CD28. All graphs represent mean +/-SD. One-way ANOVA or Student’s T-test measured statistical significance. * P<0.05; **P<0.01; ***P<0.001; ****P<0.0001.

Previously, we established a crucial role for extra-mitochondrial pyruvate metabolism in CD4^+^ T cell activation-induced enhancer remodeling^5^. To evaluate this further, peripheral blood CD4^+^ T cells obtained from healthy donors were activated by anti-CD3/CD28 stimulation for 24 hours, in the presence of synovial fluid, with or without pharmacological inhibition of pyruvate dehydrogenase (PDH). Inhibition of PDH activity resulted in a significant decrease in the expression of JIA-related genes (**Figure 5c)**. This observation, together with a lack of oligomycin inhibition, strongly supports the notion that extra-mitochondrial pyruvate metabolism plays a crucial role in histone acetylation during CD4^+^ T cell activation in the pro-inflammatory SF environment.

## Discussion

Autoimmune diseases are associated with aberrant autoreactive immune cells that lead to tissue damage resulting in increased morbidity and mortality. The intracellular metabolism of immune cell populations undergoes significant alterations in the context of autoimmunity, and concurrently, the epigenome often exhibits distinct remodelling^2,3,15–17^. While these individual phenomena are well-documented, the molecular mechanisms connecting immune cell metabolism with specific changes in the epigenetic landscape remain unclear.

Considerable evidence supports a critical role for glycolysis during T cell activation, differentiation, and function^18–24^. In the context of autoimmune disease, inhibition of glycolysis has shown promising effects, specifically in the context of rheumatoid arthritis. Studies indicate that suppressing glycolysis can reduce disease severity in autoimmune disease mouse model models^25–28^. However, there is an absence of mechanistic studies evaluating the direct connection between altered intracellular metabolism and epigenetic reprogramming in pathogenic T cells isolated directly from inflammatory environments. Additionally, for JIA, there have been no studies that have explored the metabolic profile of SF CD4^+^ T cells. Here, we demonstrate that inhibition of glycolysis during the activation of SF-derived CD4^+^ T cells is effective in preventing H3K27ac landscape remodeling (**Figure 4a-d**). Notably, this effect is particularly pronounced in H3K27ac-associated promoter regions (**Figure 4e-f**). Furthermore, the effect of inhibiting glycolysis is more robust on the promoter regions of JIA SF T cells compared to HC T cells (**Figure S4a-b**).

Increased glucose uptake plays a pivotal role in various aspects of T cell biology, including development, proliferation, and function, as demonstrated in several studies^16,24,29–32^. Notably, T cells deficient in Glut1 glucose transport exhibit impaired proliferation^30^. In this context, our data demonstrates that JIA SF CD4^+^ T cells exhibit heightened glycolytic activity and increased acetyl-CoA production compared to T cells from the peripheral blood of the same individual or healthy controls (**Figure 1d-f**). This finding contrasts with the metabolic behavior of PB CD4^+^ T cells in RA patients, which undergo a fundamental shift in glucose utilization, favoring diversion away from pyruvate and lactate production and towards the pentose phosphate pathway^33–35^. Furthermore, RA T cells have been reported to be unable to maintain the NAD+ coenzyme pool in the mitochondria, which subsequently affects the TCA cycle and aspartate production^10^. Our findings reveal a distinctive metabolic signature in T cells isolated from the SF of JIA patients. Notably, we observed no significant defects in the production of malate and fumarate metabolites within the TCA cycle when comparing JIA SF and HC T cells (**Figure 1a**). Our Seahorse experiments also indicate no substantial differences in mitochondrial respiration between JIA SF or PB T cells and HC PB T cells (**Figure 1b-c).** These observations demonstrate distinct metabolic signatures that vary between disease and CD4^+^ T cell location.

In the context of JIA, recent evidence has revealed that SF has the capacity to induce polarization towards Th1 cells, characterized as proinflammatory T helper cells^1^. Here, we demonstrate that the exposure of HC T cells to SF induces a significant increase in glycolytic activity compared to controls (**Figure 3c-d**). This increase in glycolytic response is not mirrored by an increase in mitochondrial respiration (**Figure 3a-b**), showing a selective influence of SF on the glycolytic pathway. The complexity of SF means that pinpointing a singular component responsible for induction of glycolysis in these cells is challenging. However, glycolysis-mediated alterations in the H3K27ac landscape likely play a role in the development of pro-inflammatory SF T helper cells.

Previously, we undertook a comprehensive exploration of enhancer and super-enhancer signatures using H3K27ac chromatin immunoprecipitation in CD4^+^ T cells derived from the synovial fluid of JIA patients^2^. These findings demonstrated that enhancer profiles vary significantly between subsets of T cells, particularly effector/memory cells, isolated from HC and those from JIA patients. Our current study reveals that these altered H3K27ac landscape persist even after ex*-vivo* T cell activation. We also observed that JIA-associated promotors are highly enriched for ETS1, KLF6, RUNX1, AP1 and Plagl1 transcription factor binding motifs (**Figure 2f**). These transcription factors have been previously associated with autoimmune diseases modulating the function of T cell populations^36–43^. Our observations suggest that these transcription factors may also play a crucial role in the development of pathogenic SF CD4^+^ T cells.

In our previous work we were unable to determine whether aberrant enhancer profiles are causally related to JIA disease pathogenesis or merely the consequence of the local proinflammatory environment. Here, we have evaluated the direct influence of the environment on the epigenetic landscape. Our findings reveal that SF exposure was sufficient to alter the H3K27ac landscape of HC T cells. Notably, this exposure leads to the acetylation of promoters/enhancers previously identified as active in JIA SF T cells (**Figure 3e**). As previously mentioned, we show an enrichment of JIA-associated enhancers for ETS and RUNX1 binding motifs. ETS and RUNX1 can be activated by proinflammatory signals, such as TNF-α, IL-1, and TGF-β, cytokines present in the SF^44–46^. In this way, the pro-inflammatory SF environment can promote chromatin remodeling through glycolysis-driven changes in H3K27ac, priming promoters/enhancers for subsequent cytokine-mediated transcription factor activation.

Recently we have identified nuclear pyruvate dehydrogenase (PDH), as an important glycolytic intermediate that facilitates histone acetylation and transcriptional activation after TCR-engagement^5^. The generation of pyruvate and the translocation of PDH to the nucleus was identified as a pivotal step, indispensable for producing acetyl-CoA, which, in turn, is essential for the remodeling of enhancers triggered by T cell activation. PDH has also been found to be essential for histone acetylation and subset-specific gene expression in Th17 cells^4^. In this study, we demonstrate that PDH inhibition abrogates the expression of SF-induced genes (**Figure 5b**). These findings not only emphasize the importance of PDH in this context but also support potential therapeutic interventions involving PDH inhibition for the benefit of JIA patients. Inhibition of PDH has been explored in the context of conditions such as acute myeloid leukemia, where it leads to decreased glycolysis and increased reliance on oxidative phosphorylation^47^ However, the potential application of PDH inhibitors within the autoimmune disease context remains unexplored. We propose that this approach may offer reduced toxicity and increased specificity compared to the use of glycolysis inhibitors.

Taken together, our observations suggest that the pro-inflammatory synovial environment results in metabolic dysfunction which can disrupt the epigenetic landscape resulting in transcriptional reprogramming. The dependence of activated T cells on pyruvate production through glycolysis to modulate the H3K27ac landscape and promoter/enhancer activity further supports targeting T cell glycolysis to suppress inflammatory responses and promote tolerance and immune suppression.

## Acknowledgments

We are grateful to all the authors of this paper for their contributions and significant efforts in completing and publishing this research. We would like to extend special thanks to Bas Vastert, and Jorg van Loosdregt for giving us access to patient samples and for the scientific discussions. Can Gulersonmez, Edwin Stigter, from the metabolomic facility of the UMC for their scientific discussions and for their assistance on the metabolomic studies performed on this study. Theo Chalkiadakis and Stefan Prekovic for their invaluable assistance with the RNA-seq and ChIP-seq analyses performed in this study. Finally, we would like to thank all members of the Coffer Lab for their valuable discussions concerning this work. This work is supported by grants from the Stichting WKZ Foundation (2016) and ReumaNetherlands (NR 18-01-401).

---

**Supplementary Figure 1.**
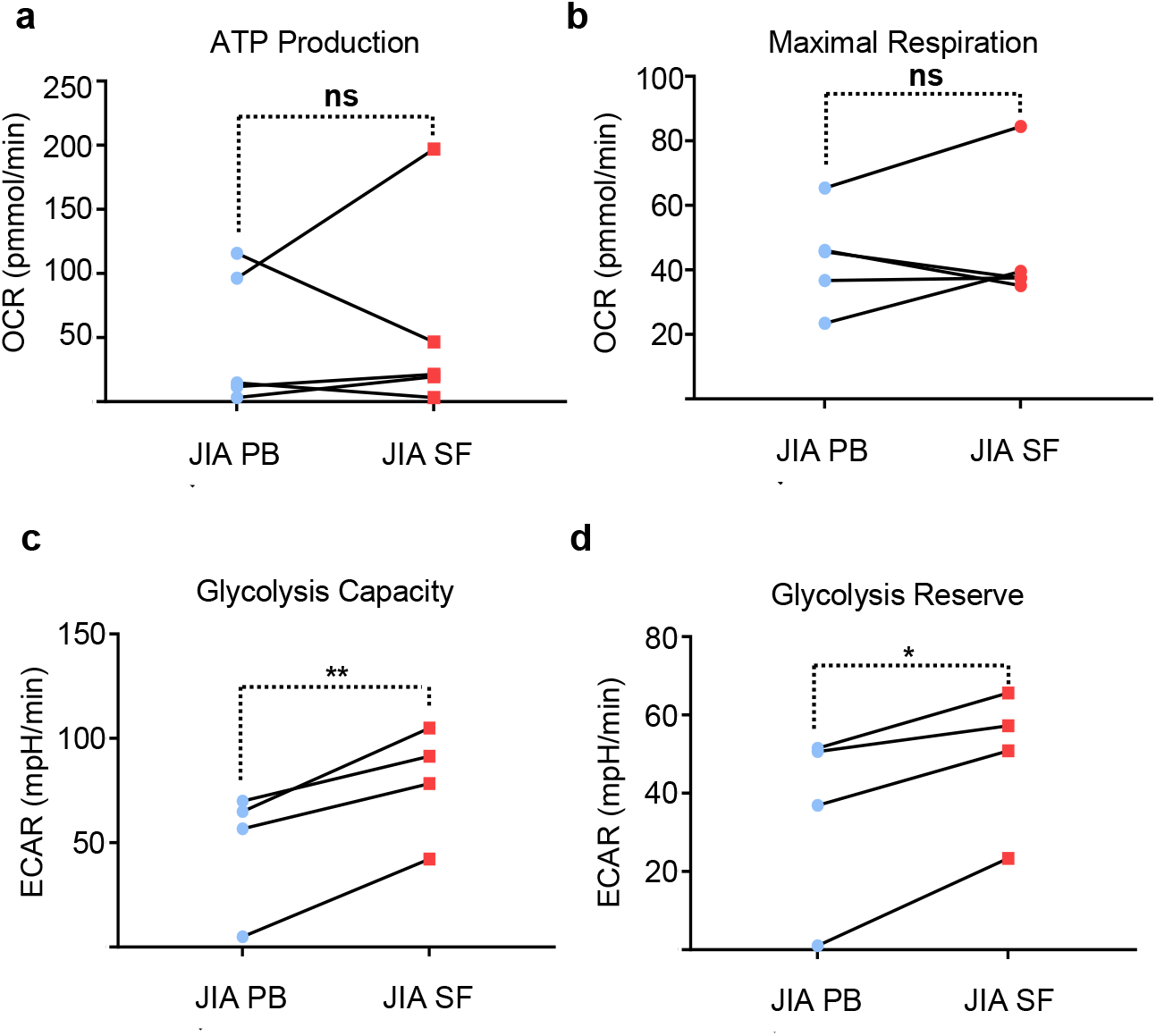
(**a-b)** JIA from PB or SF CD4^+^ T cells were activated for 24h with CD3/CD28 and extracellular acidification rates (OCR) were measured by Seahorse technology. **(c-d)** JIA from PB or SF CD4^+^ T cells were activated for 24h with CD3/CD28 and extracellular acidification rates (ECAR) were measured by Seahorse technology. All graphs represent mean +/-SD. Statistical significance was measured by Student’s T-test. * P<0.05; **P<0.01.

**Supplementary Figure 2.**
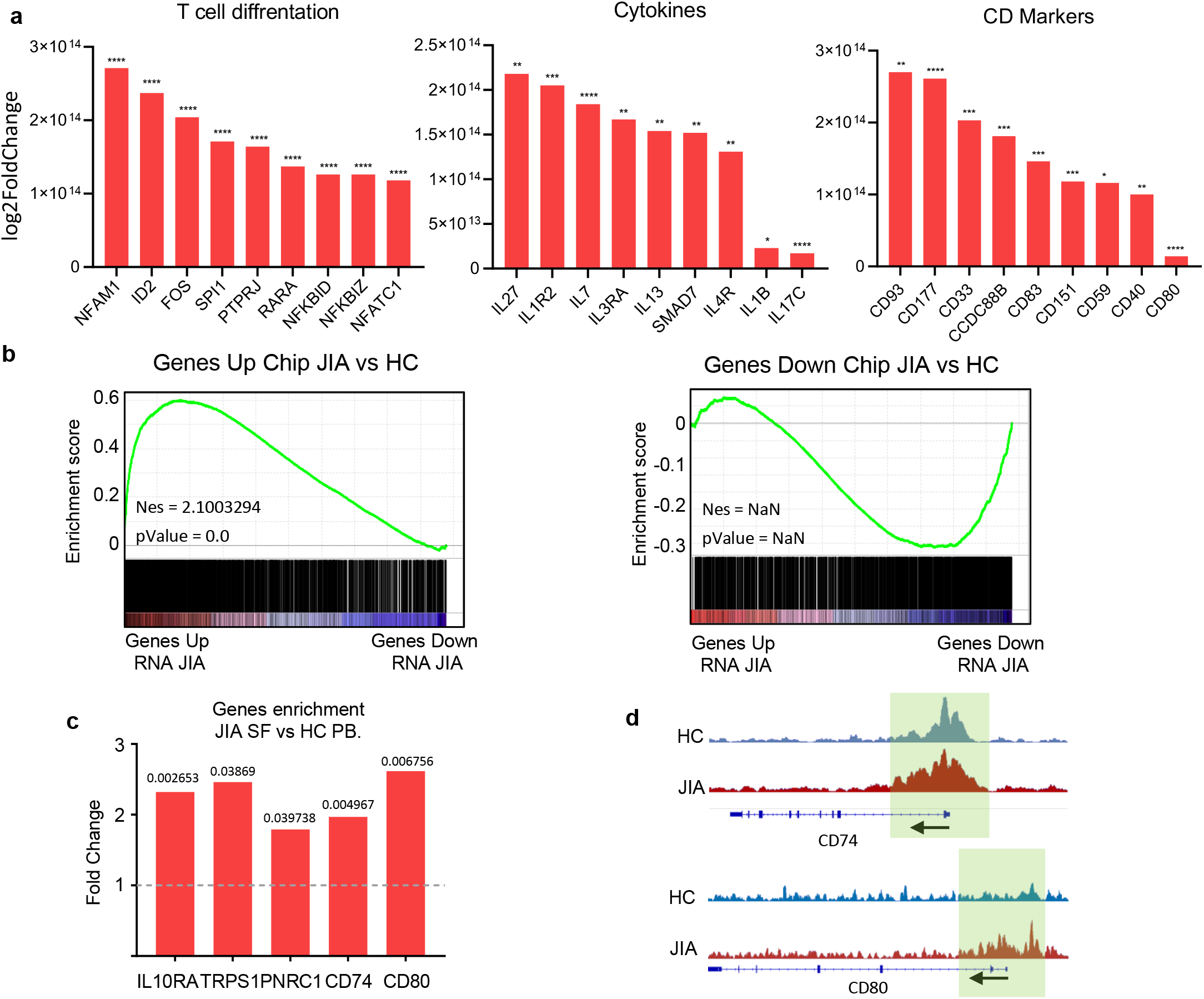
**(a)** Selected enhancer-associated genes that are upregulated in activated JIA SF-derived CD4+ T cells. **(b)** Gene set enrichment analysis between different express genes on JIA-activated CD4^+^ cells and enhancers (Es) that are up-regulated or down-regulated on JIA-activated versus HC-activated CD4^+^ T cells. **(c)** Graphical representation of genes that are up in activated JIA SF cells compare with HC T cells. **(d)** Gene tracks for CD74 and CD80 showing ChIP-seq signals for H3K27ac, on T memory cells derivate from PB HC or SF JIA. Light-green-shaded areas represent an individual enhancer. All graphs represent mean +/-SD. One-way ANOVA or Student’s T-test measured statistical significance. * P<0.05; **P<0.01; ***P<0.001; ****P<0.0001.

**Supplementary Figure 3.**
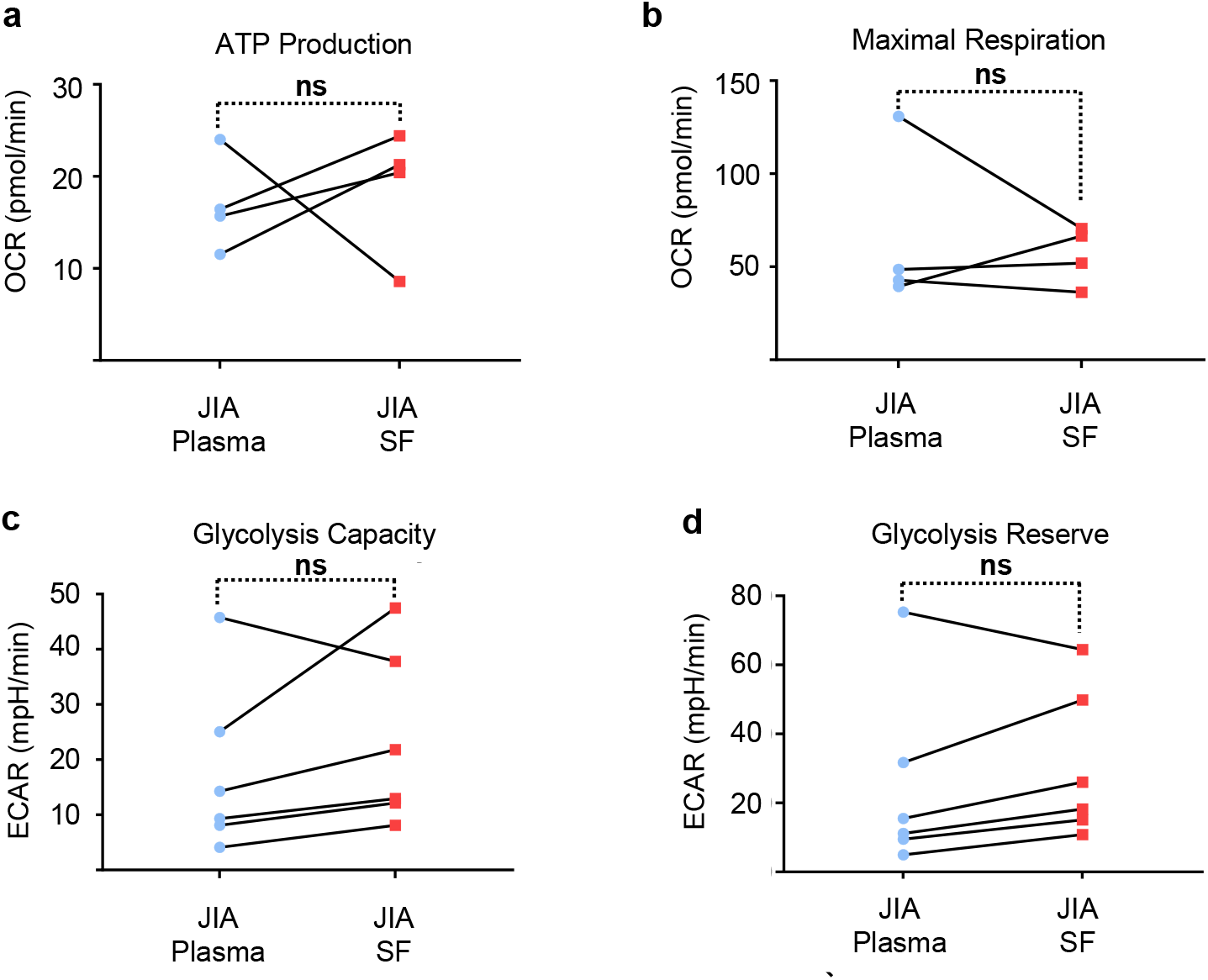
**(a-b)** HC PB CD4^+^ T cells were activated for 20h with anti-CD3/CD28 in presence of JIA SF or plasma (30%) and extracellular acidification rates (OCR) measured by Seahorse technology. **(c-d)** HC PB CD4^+^ T cells were activated for 20h with anti-CD3/CD28 in presence of JIA SF or plasma (30%) and extracellular acidification rates (ECAR) measured by Seahorse technology. All graphs represent mean +/-SD. Statistical significance was measured by Student’s T-test.

**Supplementary Figure 4.**
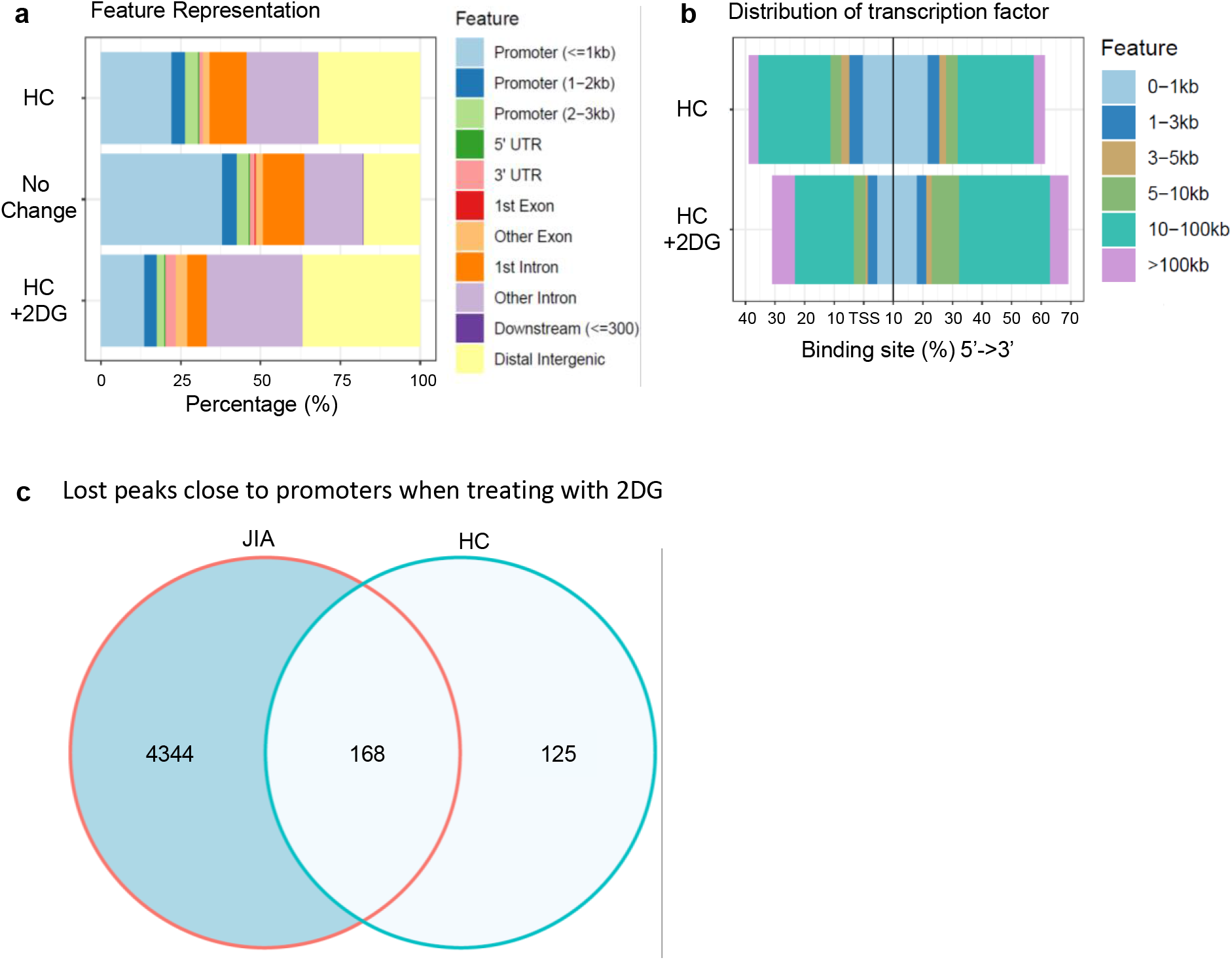
**(a)** Genomic distribution of peaks identified in ChIP-Seq data. **(b)** Distribution of transcription factor-binding loci relative to 5’ ends of genes. **(c)** A Venn diagram illustrating the intersecting peaks adjacent to promoters are lost when Hc or JIA-SF T cells are activated in the presence of 2DG.

